# Role of *BicDR* in bristle shaft construction, tracheal development, and support of *BicD* functions

**DOI:** 10.1101/2023.06.16.545245

**Authors:** Aleksandra Jejina, Yeniffer Ayala, Greco Hernández, Beat Suter

## Abstract

Cell polarization requires asymmetric localization of numerous mRNAs, proteins, and organelles. The movement of cargo towards the minus end of microtubules mostly depends on cytoplasmic dynein motors, which function as multiprotein complexes. In the dynein/dynactin/Bicaudal-D (DDB) transport machinery, Bicaudal-D (BicD) links the cargo to the motor. Here we focus on the role of *BicD-related* (*BicDR*) and its contribution to microtubule-dependent transport processes. *Drosophila BicDR* is required for the normal development of bristles and dorsal trunk tracheae. Together with *BicD,* it contributes to the organization and stability of the actin cytoskeleton in the not-yet-chitinized bristle shaft and the localization of Spn-F and Rab6 at the distal tip. We show that *BicDR* supports the function of *BicD* in bristle development and our results suggest that BicDR transports cargo more locally whereas BicD is more responsible for delivering functional cargo over the long distance to the distal tip. We identified the proteins that interact with BicDR and appear to be BicDR cargo in embryonic tissues. For one of them, EF1γ, we showed that *EF1γ* genetically interacts with BicD and *BicDR* in the construction of the bristles.

## Introduction

Microtubules are crucial for the growth of polarized cells. At the same time, they attach different cellular organelles, such as the nucleus, the Golgi apparatus, and the endoplasmic reticulum, to a specific cellular compartment and enable the polarized transport of vesicles, mitochondria, mRNAs, and cytoskeletal elements^1,2^. Because of their growth, which is focused towards one pole of the cell^3^, the *Drosophila* macrochaetae can serve as a model tissue for studying such cytoskeleton-dependent transport processes that are necessary for bristle development^4^. Several studies indicate that endocytic trafficking has an important function in process^5^. Multiple defects in bristle development have been described in flies that are mutant for members of the *Rab* gene family, which are known to regulate intracellular vesicle trafficking. Whereas *Rab6* and *Rab11* mutants eclose with short and stubble-like bristles, *Rab35* mutants display forks and kinks in their macrochaetae^6–9^.

The *Drosophila* Bicaudal-D (BicD) is part of an evolutionarily conserved transport machinery, the microtubule-dependent dynein/dynactin transport apparatus. Its essential functions in the development of the oocyte and embryo are well characterized^10–13^. Furthermore, it was observed that *BicD^A40V, S103^* and *BicD^null^* mutants display short sternopleural and scutellar bristles^14^. This mutant phenotype pointed to a function of *BicD* in the development of macrochaetae, one that had not been studied so far. The similarity between the bristle phenotype of *BicD* and *Rab6* suggested possible interactions between these two in constructing the bristles. Support for this hypothesis comes also from work that showed that Rab6 and BicD function together in the delivery of secretory pathway components^15^.

The BicD protein family contains another member, BicD-related (BicDR, enoded by *CG32137*), which was discovered due to its strong sequence homology to BicD. In *Danio rerio,* BicDR is needed for the pericentrosomal transport of Rab6-positive vesicles during neural development. To perform this function, the *Danio rerio* BicDR requires the lysine residue K512, which is highly conserved between BicD and BicDR paralogues and orthologues^10^. The *Drosophila* BicD sequence around K730 is essential for the interaction with the cargo^16^ and a point mutation in this codon resulted in the isolation of the first single amino acid substitution that produced a *BicD^null^* phenotype, indicating that this lysine is key to the physiological role of BicD^17^. The homologous lysine in Drosophila BicDR is conserved and located at position K555 of BicDR-B and K461 of BicDR-A. For simplicity, we will refer to this residue as K555 for both isoforms. However, whether K555 of *Drosophila* BicDR serves the same function as its K730 orthologue of BicD remains to be tested. There are also interesting differences between BicD and BicDR. While *Drosophila* BicD consists of three coiled-coil domains, only the first and third are conserved in the BicDR protein.

Although the strong homology between the fly BicD and BicDR suggests similar functions, the role of *BicDR* in *Drosophila* has not yet been examined. The homology is strongest in the coiled-coil domain near the C-terminus which, in the case of BicD, is known to be needed for the attachment and transport of various cargoes^10,16^.

We set out to investigate the function of *BicDR* with a focus on a potential role in MT-dependent trafficking and possible cooperation or competition between BicDR and BicD that might contribute to the development and maintenance of polarized cell growth. Here we describe the genetic interaction between *BicD* and *BicDR* and its contribution to fly development. Furthermore, we describe and compare the effect of different *BicD* and *BicDR* alleles on the formation of the macrochaetae and trachea.

## Materials and Methods

### Fly stocks and genetics

Flies were kept and bred on standard cornmeal agar containing yeast, sucrose, potassium sodium tartrate, methylparaben, and propionic acid. For the crosses, multiple virgins (5 to 10) were added to several males (3 to 5) and incubated at 25°C. The following fly strains were used in this study: *BicD^PA66^*, *BicDR^(rev)^*, *BicDR^29^*, *BicDR^51^*, and *BicDR^71^*. Standard methods were used to generate *BicDR* excision stocks with the two *P* elements *P{SUPor-P} and P{RS5}*^18,19^. The excisions were characterized molecularly by extracting DNA from heterozygous mutant males followed by PCR with primers framing the deleted regions of the *BicDR* gene. The screening by PCR revealed that two excision stocks are missing a fragment around the insertion site of the P-elements *P{SUPor-P}*: *BicDR^29^*, *BicDR^51^*, and one around the insertion site of *P{RS5}*: *BicDR^71^*. These stocks were double-balanced and kept for further examinations. In the case of the mutant *BicDR^(rev)^*, the activated P-element *P{RS5}* reverted the genomic sequence of the *BicDR* gene to the wild-type sequence when it jumped out. The wild-type revertant *BicDR^(rev)^* was used as a control for the excision mutants.

*v; CyO/Sp* flies were kindly provided by Simon Bullock. Stocks from the Vienna Drosophila RNAi Center (VDRC) and the Bloomington Stock Center are listed in the appendix. For tissue-specific knock-down or gene expression, the *UAS-Gal4* system was used^20^.

### CRISPR/Cas9 and generation of transgenic flies

All gRNAs were designed manually and verified on the web-based tool called *CRISPR optimal target finder*^21^. The gRNAs for attaching a GFP-tag (5’-ATTATCGCTGAAATAAACTC-3’) and the gRNA for the deletion and substitution of K555 (5’-AGTCCATTCAGCAAAAGG-3’) were cloned into pCFD5 plasmids following the “gRNA cloning protocol for cloning single gRNA plasmids”^22^. Transgenic flies were generated using the ΦC31-based integration system^23^ and crossed with *nos-Cas9* expressing animals.

To add a GFP-tag to the C-terminus of the BicDR protein, the appropriate *eGFP* DNA sequence^24^ with a linker and two 1,200 bp long arms homologous to the *BicDR* gene and framing the stop codon were cloned into a pBluescript II SK (+) vector. The construct was injected into embryos with the genotype w, y, w^+^ *nos-Cas9*/Y; gRNA *v^+^*/+; *BicDR\**/*BicDR\**. To generate the *BicDR^K555A^::GFP* mutant, the sequence within the template vector was modified by site-directed mutagenesis before injection. All constructs were sequence verified. All primers used for DNA construction are listed in the appendix.

### Genetic interaction assay

In the following crossing schemes *BicDR\** indicates one of the excisions, *BicDR^29^*, *BicDR^51^*, and *BicDR^71^*, the deletion mutant *BicDR^8.1^*, the *BicDR^null^* allele *Df737* (*BicDR^Df^*) or the revertant *BicDR^(rev)^*.

♀ w; *BicD^PA66^*/CyO; *BicDR\** / TM6B x w; *Df7068*/CyO; *Df4515/*TM3, Sb ♂

For every cross, 30 virgins with the above genotype were added to 15 males. Every two days, the flies were transferred to a new plastic bottle. The frequency of every genotype of the progeny was determined. The progeny was also sorted by sex and genotype and kept at 18°C for the following experiments. Females that were not virgins anymore were dissected, and their ovaries were stained. The frequency of genotypes of eclosed flies from each cross was counted for 9 days each. All statistics and graphics were done using the GraphPad Prism 5 software.

### Analysis of bristle development

The following crosses were used to determine the severity of the *BicDR* alleles:

♀ w; *BicD^PA66^*/CyO; *BicDR\** / TM6B x *Df7068*/SM6B ♂ → w; *BicD^PA66^*/*Df7068*; *BicDR\** /+

♀ w; *BicD^PA66^*/CyO ftz lacZ x w; *Df7068*/CyO; *Df4515, w^+^/*TM3, Sb ♂ → w; *BicD^PA66^*/*Df7068*; *Df4515, w^+^*/+

To investigate the bristle phenotype further, pupae with the following genotypes were dissected and stained at their appropriate stage following the protocol by Tilney et al.^25^

♀ w; *BicD^PA66^*/ CyO, Act-GFP; *BicDR\**/TM6B, Tb x Df7068/CyO, Act-GFP ♂

→ w; *BicD^PA66^*/*Df7068*; *BicDR\**/+

→ w; *BicD^PA66^*/ *Df7068*; TM6B, Tb/+

### Analysis of tracheal development

The following stocks and crosses were used to study the development of the trachea:

→ w; *BicDR^71^*/ *BicDR^71^*

♀ w; *BicD^PA66^*/ CyO, Act-GFP; *BicDR^71^*/ *BicDR^71^* x w; *BicDR^71^*/ *BicDR^71^* ♂

→ w; *BicD^PA66^*/ +; *BicDR^71^*/ *BicDR^71^*

### Immunostaining and microscopy

Dechorionated embryos or dissected tissue that was kept on ice for less than 30 minutes were fixed in 4% paraformaldehyde for 20 minutes and blocked with either 5% milk or bovine serum albumin (Fraction V) for 2 hours at room temperature. Primary antibodies were incubated overnight, followed by washing steps and incubation with secondary antibodies for at least 2 hours. Primary antibodies were diluted as follows: anti-GFP (rabbit, 1:200, ImmunoKontact), anti-GFP (mouse, 1:200), anti-Serp (rabbit, 1:300)^26^, anti-Ef1γ (rat, 1:1,000)^27^, anti-Rab6 (rabbit and guinea pig, 1:200)^28^, anti-Spn-F (rabbit, 1:300, DSHB), anti-Asense (guinea pig, 1:100)^29^. Secondary antibodies were Alexa Fluor 488 (anti-rabbit 1:800), Alexa Fluor 488 Plus (anti-rabbit, 1:200), Alexa Fluor 647 (anti-mouse, 1:200), Alexa Fluor 647 Plus (anti-rabbit, 1:200), Cy3 (anti-mouse, 1:400). The images were taken with a Leica TCS-SP8 confocal laser-scanning microscope and processed using the FIJI software.

### Isolation of embryonic *BicDR* complexes for mass spectrometry

12-16 hours old embryos were collected and lysed in homogenization buffer (25 mM Hepes pH 7.4; 150mM NaCl; 0.5mM EDTA pH 8.0; 1 mM DTT and 1 tablet of proteinase inhibitor cocktail; Roche 11836170001). The aqueous phase of the lysate was collected after 1 hour of centrifugation at 4°C and centrifuged again for 25 minutes. Subsequently, one part of the aqueous phase was saved as input control, while the rest was incubated with Plus Sepharose G beads that were coated with anti-GFP antibody overnight at 4°C. 5-7 washing steps with wash buffer (25 mM Hepes pH 7.4; 150mM NaCl; 0.5mM EDTA pH 8.0; 1 mM DTT and ½ tablet of proteinase inhibitor cocktail, Roche 11836170001) were performed before the beads were either sent for mass spectrometry or prepared with the appropriate amount of SDS for SDS/PAGE and Western blot. SDS/PAGE bands that were present in the IP from the BicDR::GFP-fusion protein were cut out of the gel and sent directly for mass spectrometry. As a control, the equivalent regions of the control lanes were also cut out and used for a mass spectrometric analysis.

### Yeast two-hybrid assay

The full-length cDNA of *BicDR* and *Ef1γ* as well as the C-terminal domain (CTD) of *BicDR* were cloned into pOAD and pOBD2 vectors so that they were in frame with the activator domain (AD) or the DNA binding domain (BD)^13,30^. In this way, BicDR-AD, BicDR-CTD-AD, Ef1γ-AD, as well as BicDR-BD, BicDR-CTD-BD, and Ef1γ-BD were created. The BicD-AD and Egl-AD, as well as BicD-BD and Egl-BD, have been described previously^13^.

## Results

### Functional redundancy between *BicDR* and *BicD*

While BicD consists of three coiled-coil domains, only the first and third are conserved in the related BicDR protein (Figure S1A). The homology is strongest in the coiled-coil domain near the C-terminus, which in the case of BicD is known to be needed for the transport of different cargoes^16,31^. Although the N-terminus is less highly conserved between the two proteins, it was shown that BicD and BicDR require it for the efficient binding to dynein and dynactin *in vitro*^32,33^.

The *Drosophila BicDR* extends over 24.4 kb of genomic sequence with 6 exons in total. Remarkable is the relatively long intron with about 18 kb between the first and second exons, which is also similar to the structure of the *BicD* gene^34^. There are two transcripts: *BicDR-A* and *BicDR-B* with the difference that the *BicDR-A* start codon is localized 282 bp downstream of the *BicDR-B* one (Figure S1B). Both transcripts share the reading frame and the stop codon. The extra peptide of BicDR-B contains a repeat of 5 Asparagine and 6 Serine residues, respectively, but shows no homology to any other gene or organism^35^.

To identify specific *BicDR* alleles that could be null alleles, we picked two P-element insertions and created imprecise excision mutations (see Methods section). For further genetic analyses, we retained two excision lines from the upstream element and one from the downstream insert: *BicDR^29^*, *BicDR^51^*, and *BicDR^71^* (Figure S1B). While the excision *BicDR^29^* removed only the 5’ UTR region, excision *BicDR^51^* removed in addition to that also the entire first protein-coding exon. *BicDR^71^* is the only excision that removes the second, third, and fourth protein-coding exon and thereby also induces a stop codon in the first coiled-coil domain of BicDR-A and BicDR-B (Figure S1). In addition, we retained a precise excision *BicDR^(rev)^* from the downstream element as a control. We also generated a precise mutation where the Q554 and K555 codons in *BicDR* were deleted (*BicDR^8.1^*, see Figure S1B).

All the described *BicDR* mutants were viable and fertile, indicating that *BicDR* is a non-essential gene. However, hemizygous *BicDR^71^* females eclosed with individual macrochaetae that contained discolored and brittle tips that broke off easily (Figure 1; white arrow pointing to a white bristle tip in Figure 1C). Additionally, a fraction of the female adults contained additional aSC, aPA, or pNP macrochaetae^36^ (see white arrows pointing to shorter pSCs and a blue arrow to an additional aSC in Figure 1D). This phenotype was not observed in *BicDR^29^* mutants. Knocking down *BicDR* by RNAi driven with the *en-Gal4* driver also led to adult females eclosing with individual slightly shorter aSC or pSC macrochaetae (Figure 1E). This phenotype could be observed significantly more often in females than in males. 12 out of 18 female and 2 out of 8 male flies eclosed with at least one shorter bristle, while no control animals (0 out of 32 *en-Gal4; UAS-GFP*) displayed such a phenotype (Figure 1F). These results show that *BicDR* functions in the formation and development of mechanosensory organs of *Drosophila* during metamorphosis.

**Figure 1.**
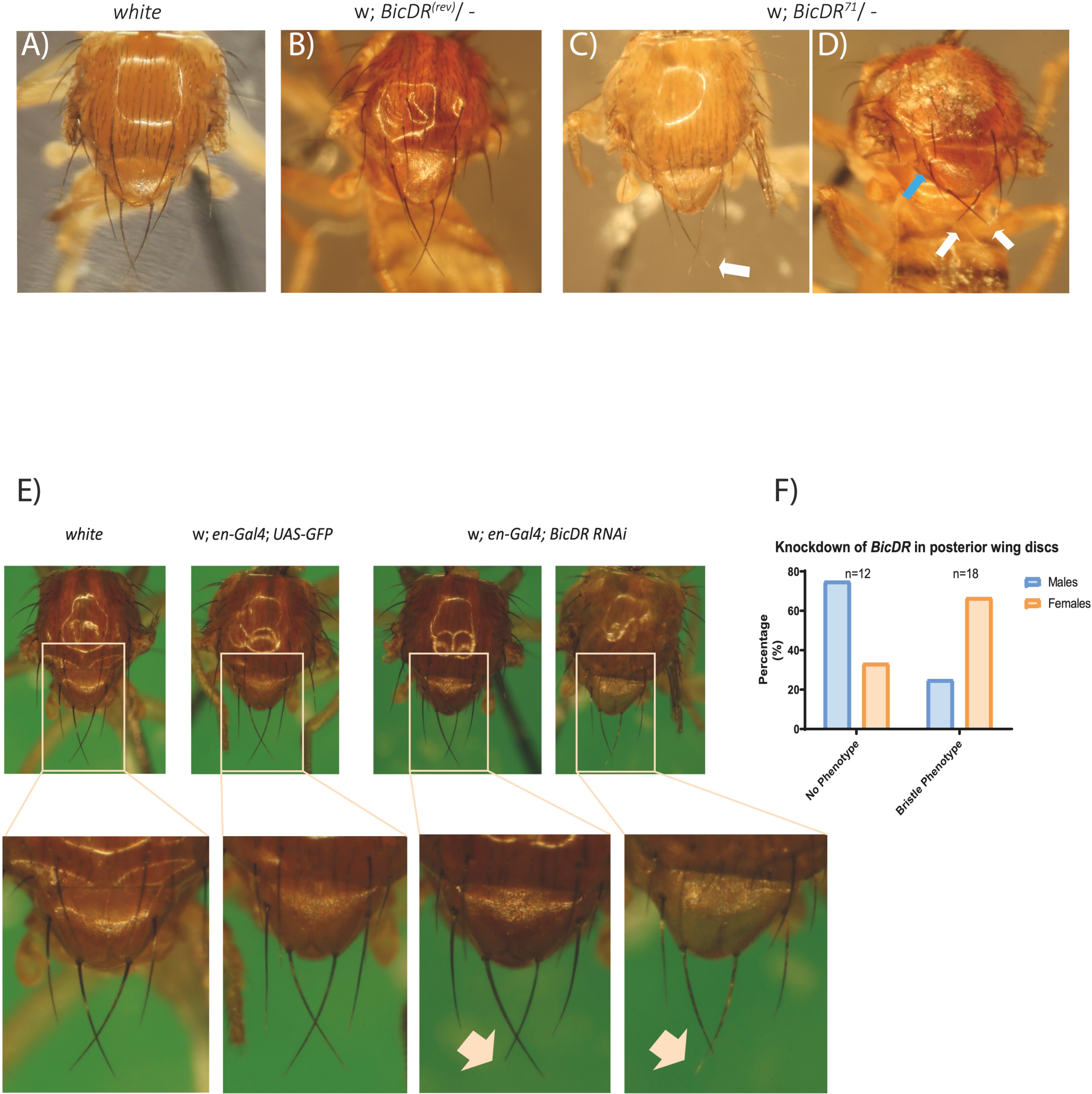
Bristle phenotypes in *BiCDR* mutants. **(A)** Macrochaetae of a wild-type control fly, the revertant w; *BicDR^(rev)^*/*Df4515* **(B)** and w; *BicDR^71^*/*Df4515* **(C and D)**. Note that the notum of the w; *BicDR^(rev)^*/*Df4515* control does not show any differences compared to the wild type, whereas w; *BicDR^71^*/*Df4515* flies eclosed with shorter pSC macrochaetae (white arrows in **D**) and occasionally with an additional aSC bristle (blue arrow in **D**); identification of macrochaetae was according to Ref^36^). **(E)** RNAi knockdown of *BicDR* induces defective bristles. w; UAS-BicDR-RNAi/en-Gal4; UAS-BicDR-RNAi/UAS-GFP. Note slightly paler macrochaetae tips and occasional shorter macrochaetae marked with an arrow. Mostly female flies eclosed with this knock-down phenotype of *BicDR* (12 out of 18 female and 2 out of 8 male flies). **(F)** Comparison with wild type and w; en-Gal4/ CyO; UAS-GFP controls.

The sequence similarity between BicD and BicDR suggests that the two proteins might either be functionally redundant or compete with one another. To test these possibilities we produced flies that simultaneously carry mutations in both genes using a female sterile allele of *BicD* (*BicD^PA66^*)^37,38^. *BicD; BicDR* double mutants of the genotype w; *BicD^PA66^*/−; *BicDR\**/− were tested for viability and fertility. *BicDR\** stands for the different *BicDR* alleles tested (*BicDR^29^*, *BicDR^51^*, *BicDR^71^*, and *BicDR^8.1^*) and the wild-type revertant *BicD^(rev)^* that served as a control.

As shown in Figure 2A, hemizygous double mutant males and females were virtually absent from the offspring but appeared in the control w; *BicD^PA66^*/−; *BicDR^(rev)^*/− (13% of the total number of eclosed progeny, which is the expected frequency for the control). The genotype w; *BicD^PA66^*/−; *BicDR^71^*/− was only found in three male flies (0.65% of all eclosed progeny; Figures 3A). No progeny of the genotypes w; *BicD^PA66^*/−; *BicDR^51^*/− and w; *BicD^PA66^*/−; *BicDR^Df^*/− or w; *BicD^PA66^*/−; *BicDR^8.1^*/− eclosed, while 21 animals with the genotype w; *BicD^PA66^*/−; *BicDR^29^*/− eclosed (5.37% of the total eclosed progeny). The genotype *BicD^PA66^*/− is viable, but if both copies of *BicDR* are additionally null or strong loss-of-function alleles, the flies are not viable anymore. We conclude that one functional copy of *BicDR* is sufficient to support the residual *BicD* function in *BicD^PA66^*/− and maintain viability. This points to a redundant role of *BicD* and *BicDR* for an essential function.

**Figure 2.**
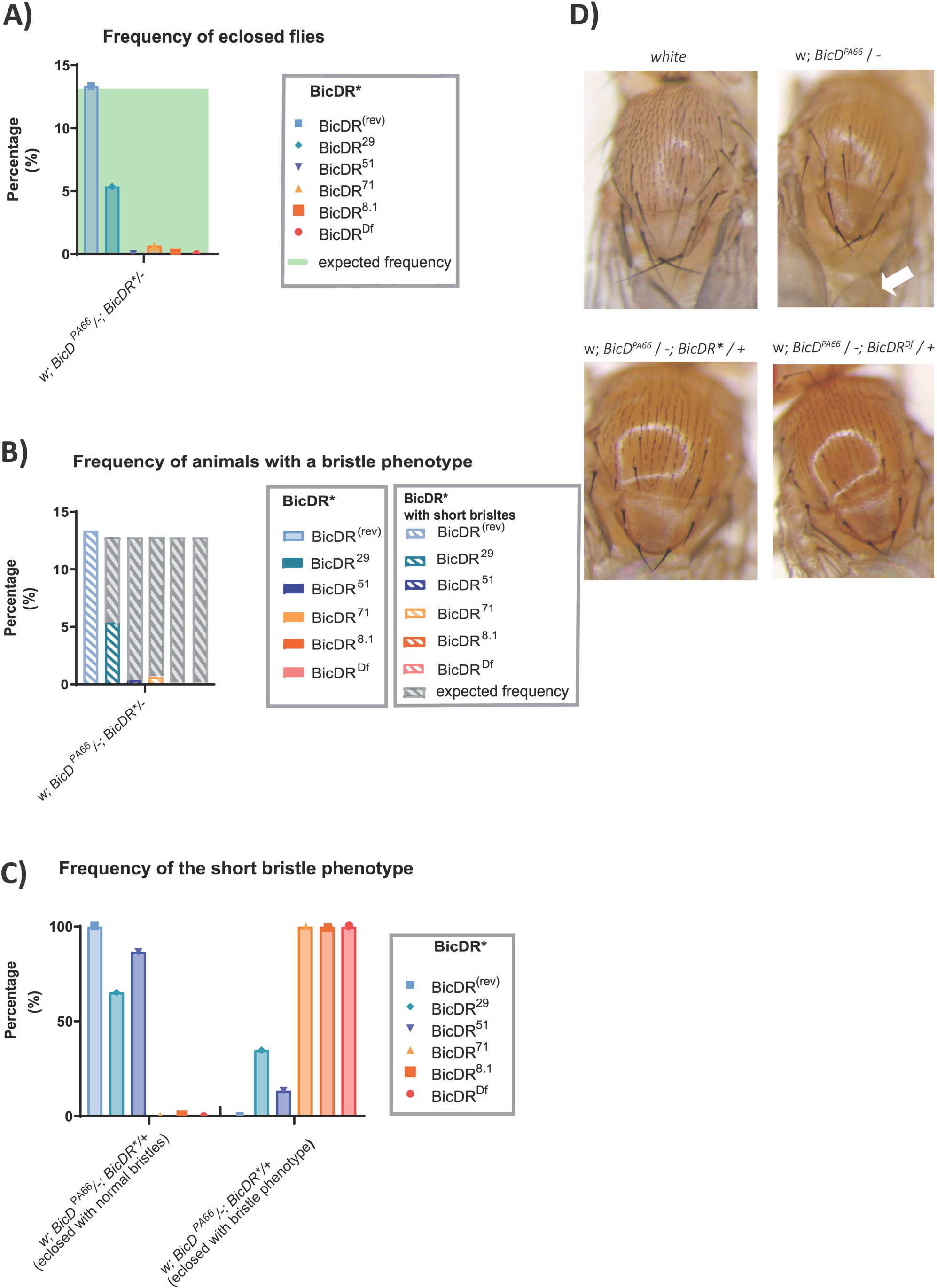
(A) Frequency of eclosed *BicD; BicDR* double mutants with the genotype w; *BicD^PA66^*/−; *BicDR*/−*. Adult flies eclosed only in crosses containing *BicDR^29^* or the control *BicDR^(rev)^* except for one escaper with the allele *BicDR^71^*. All of them developed a short-bristle phenotype. The genotype w; *BicD^PA66^* /−; *Df4515*/+ eclosed in all crosses and all flies with the named genotype developed the short-bristle phenotype as well**. (B)** shows the frequency of the short bristle phenotype in *BicD^PA66^/−; BicDR*/−* animals and **(C)** the frequency of the short bristle phenotype in *BicD^PA66^/−; BicDR*/+* animals. The *BicDR*-excisions that the animals carry and the deficiency *Df737* (this deficiency is also referred to as *BicDR^Df^*), are indicated. The calculated expected frequency is always shown in green **(A)** or gray **(B)**. **(D)** Comparison of the bristle phenotypes observed in controls (*white*), *BicD^PA66^*/−, *BicD^PA66^*/−; *BicDR^71^*/+, and *BicD^PA66^/−; BicDR^Df^/+*. Note that the *BicD* bristle phenotype, which manifests itself in discolored and brittle bristle tips, deteriorates with the reduction of *BicDR* function. The phenotypes observed in *BicD^PA66^*/−; *BicDR^71^*/+ and *BicD^PA66^*/−; *BicDR^Df^/+* show the same severity, indicating that the allele *BicDR^71^* behaves like a *BicDR^null^* mutant.

**Figure 3.**
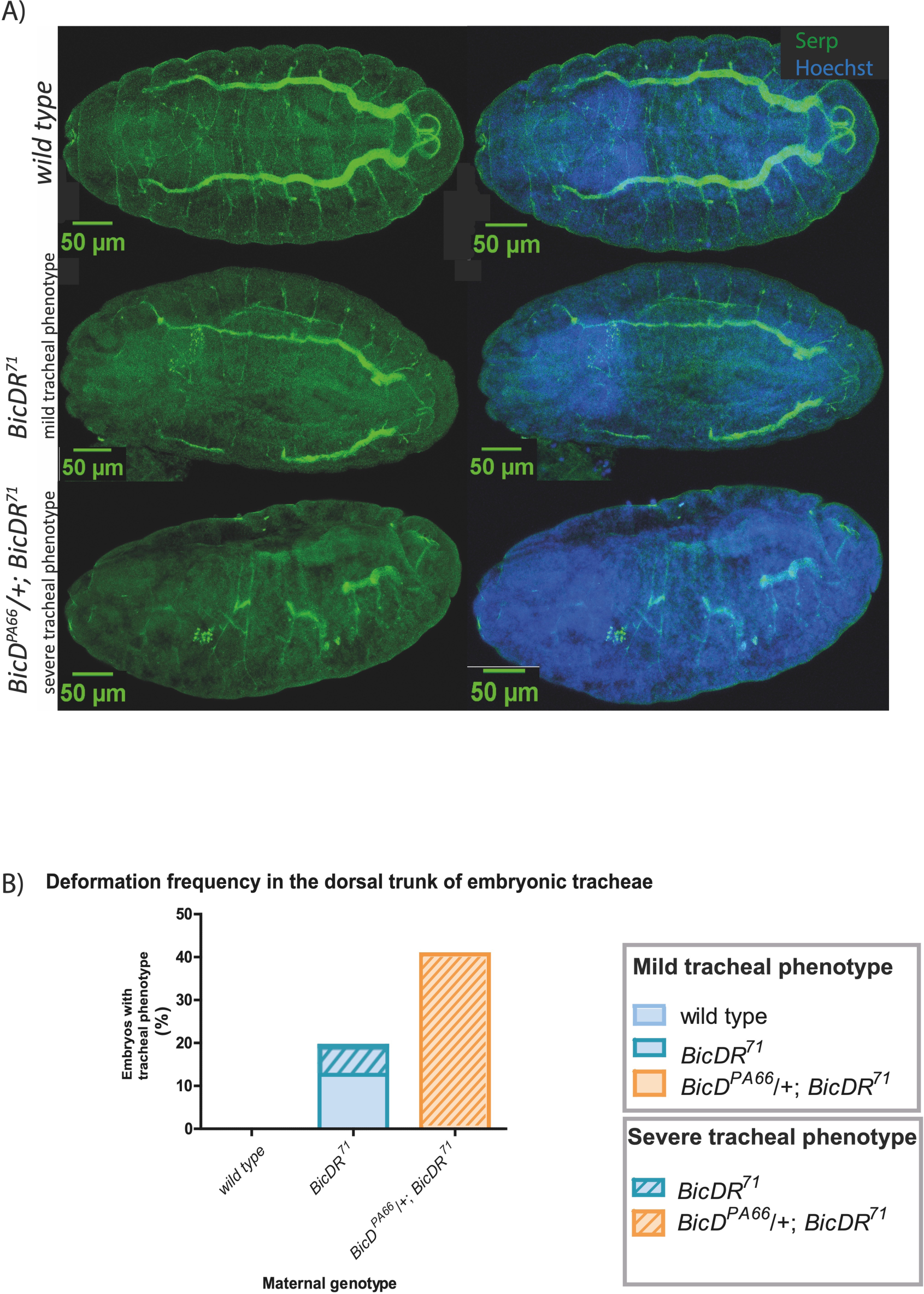
(A) Anti-Serpentine staining of late stage 13 to stage 15 embryos from wild type (*w*), homozygous *BicDR^71^*, and homozygous *BicDR^71^* mutants with a heterozygous mutation in *BicD* (*BicD^PA66^*). Serpentine is shown in green, and Hoechst is in blue. Homozygous *BicDR^71^* mutants develop mostly single gaps in one of the dorsal tracheal trunks (mild tracheal phenotype) while w; *BicD^PA66^*/+; *BicDR^71^* misses to form a dorsal tracheal trunk completely in some of the cases (severe tracheal phenotype). The frequency of the observed phenotypes is shown in **(B)**, where over 30 embryos of each genotype had been analyzed. The occurrence of malformations in the dorsal trunk in *white* embryos is 0%, in *BicDR^71^* embryos 19.3%, out of which 67% displayed a mild and 33% a severe tracheal phenotype. In w; *BicD^PA66^*/+; *BicDR^71^* embryos 40.63% of all analyzed embryos displayed the severe tracheal phenotype, while no milder phenotypes could be observed.

The few *BicD^null^* animals that survived to adulthood displayed a bristle defect phenotype with colorless and brittle bristle tips. Whereas the discolored tips were already seen in *BicD^PA66^*/− flies (Figure 2D), the shorter bristles only appeared when *BicDR* activity was also reduced in this background (*BicD^PA66^*/*-; BicDR\**/+). These animals eclosed with significantly shorter, stubble-like macrochaetae (Figure 2D). This short bristle phenotype was the strongest in flies that carried the *BicDR^71^* and *BicDR^8.1^* allele: all adult progeny with a hemizygous copy of *BicD^PA66^* and one *BicDR^71 (or 8.1)^*/+ chromosome showed the short bristle phenotype (Figure 2C). The same was true when the *BicDR* deficiency chromosome (*BicDR^Df^*) was tested in the same way (*BicD^PA66^*/*-; BicDR^Df^*/+). In contrast, less than half of the hemizygous *BicD^PA66^* flies containing *BicDR^51^* and *BicDR^29^* eclosed with short bristles. For *BicDR^29^*, these were 35% (8 out of 23 flies) and for *BicDR^51^* 33% (5 out of 15 flies). These results show again that *BicDR^+^*supports *BicD^PA66^* in bristle development but only a single functional copy of *BicDR* is not sufficient to allow normal bristle development.

The few eclosed double mutants with the genotype *BicD^PA66^*/−; *BicDR\**/− all display the described short bristle phenotype (Figure 2B). This includes the control *BicDR^(rev)^* because its third chromosome deficiency removes the *BicDR* gene completely.

The molecular and genetic analyses of the mutant *BicDR* alleles define an allelic series. The *BicDR^29^* is a hypomorphic allele and produces the weakest phenotype because hemizygous *BicD^PA66^* animals that are also hemizygous for *BicDR^29^* are viable, whilst the analogous genotype is lethal for *BicDR^71^*, *BicDR^51^*, or *BicDR^Df^*. *BicDR^29^* may have retained the most *BicDR* activity because this excision only removed the 5’ UTR region and intron sequences of *BicDR* and none of the protein-coding regions (Figure S1). We note that it still retains considerable functional activity. On the other hand, the *BicDR^71^* allele is the strongest. Our results reveal that the allele *BicDR^71^* induces the strongest effect within flies that contain the hypomorphic mutation *BicD^PA66^*. We can further conclude that *BicDR^71^* is a stronger *BicDR* allele than *BicDR^51^* and that its behavior can be compared to the deficiency of *BicDR*, *BicDR^Df^*, which removes the *BicDR* gene completely. *BicDR^71^* removes exons 2 and 3, while *BicDR^51^* removes the 5ʹ UTR region and the exon 1 of *BicDR-A* and *-B* (Figure S1). Accordingly, the results reveal an allelic series from the weakest to the strongest *BicDR* allele (*BicDR^29^< BicDR^51^<BicDR^71^* and *BicDR^Df^*).

One mechanism by which BicDR might support BicD function is suggested by their similar structure. BicD functions as a dimer, which it forms through its coiled-coil domains. If the homologous coiled-coil domains interact, BicDR might replace a BicD subunit in the active complex. To test whether BicD and BicDR form heterodimers, we performed immunoprecipitations with embryos expressing GFP-tagged BicDR and we tested for this interaction in a yeast two-hybrid experiment, which would reveal direct interactions between the two proteins. The yeast two-hybrid experiment confirmed that BicDR forms homodimers (Figure S2), as had been already described^33,39^. However, the experiment did not reveal any direct interaction between BicD and BicDR. Similarly, the embryonic immunopurification did not show copurification of the two related proteins. We thus conclude that BicDR is unlikely to support BicD function by forming dimers with BicD through their related coiled-coil domains.

### Function in tracheae development

According to the FlyAtlas Anatomical Expression Data^40^, the expression of *BicDR* is described as “very high” in the larval trachea. To examine the potential function of *BicDR* in the development of tracheae, maternal and zygotic homozygous *BicDR^71^* mutants as well as selected zygotic progeny, which were heterozygous for *BicD^PA66^* on the second chromosome and also homozygous for *BicDR^71^* on their third chromosome, were collected and stained with an anti-Serpentine (anti-Serp) antibody that marks the lumen of trachea from late stage 13 on^26^. As shown in Figure 3A, the staining of *BicDR^71^* embryos revealed that the *BicDR* mutants retain single gaps within the dorsal trunk of the tracheae, while the double mutants *BicDR^PA66^*/+; *BicDR^71^*/*BicDR^71^* showed prominent interruptions in the dorsal trunk with isolated tracheal metameres that were unable to connect through branch fusion. However, not all mutant embryos developed such tracheal malformations. To determine the penetrance of this phenotype, more than 30 embryos per genotype were examined for abnormalities in tracheal fusions. The quantification showed that 19% of maternal and zygotic homozygous *BicDR^71^* mutants (6 out of 31 embryos) and 41% of maternal and zygotic w; *BicDR^PA66^*/+; *BicDR^71^*/*BicDR^71^* embryos (13 out of 32 embryos) displayed a phenotype similar to the one shown for the respective genotype in Figure 3A, while the *white* control showed 0% of tracheal phenotypes (0 out of 35 embryos; Figures 3A, 4B).

**Figure 4.**
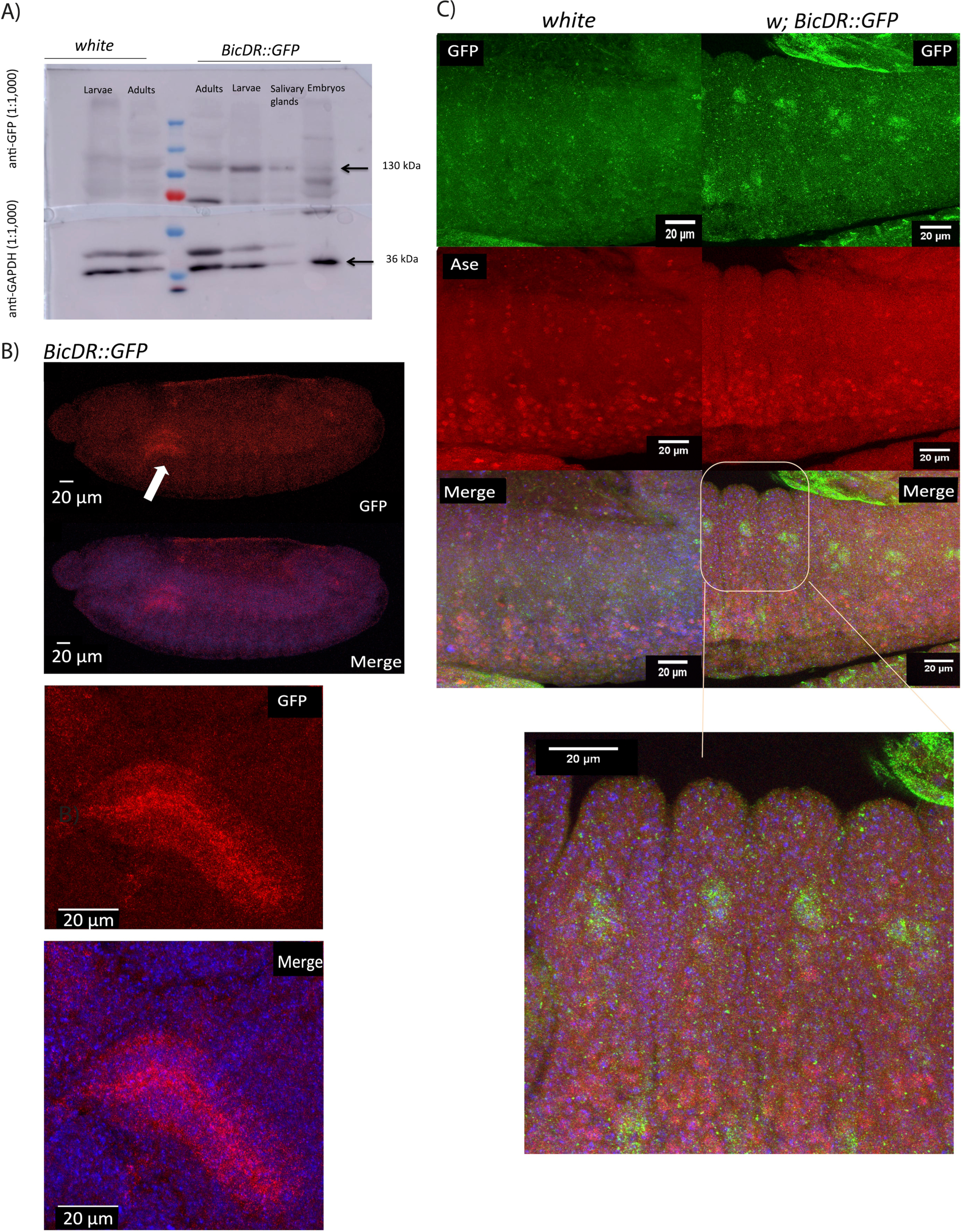
BicDR is expressed in the region of SOPs of stage 13 old embryos. **(A)** Expression during the different stages of the life cycle is shown by a Western blot stained for GFP to reveal the expression of the endogenously tagged *BicDR*. The samples loaded were stage 13 to 16 embryos, 3. instar larvae, adult flies, and salivary glands. All were of the genotype w; *BicDR::GFP*/*Df4515* or the negative control (*white*). The loading control was GAPDH with a size of 35 kDa (see lower blot). BicDR-B::GFP, with a size of 130 kDa was found mostly in adult flies, 3rd instar larvae, and the dissected salivary glands of the 3rd instar larvae, while BicDR-A::GFP with 110 kDa was mostly expressed in late embryos. **(B)** Stage 13 embryos stained for BicDR::GFP (green). The DNA is stained with Hoechst (blue). BicDR::GFP is expressed apically in the cells of salivary glands and cells along the anterior-posterior embryo axis in a metameric pattern. **(C)** Co-staining of BicDR::GFP embryos with the SOP marker Asense (red) and GFP (green) identifies the GFP^+^ cells in the vicinity of elevated Ase staining.

### BicDR::GFP is expressed in the salivary glands and in the embryo in a metameric pattern

To determine in what tissues and during which embryonic stages BicDR is expressed, we tagged the *BicDR* gene endogenously with GFP using CRISPR/Cas9 and immunolocalized BicDR::GFP in embryos after fixation. This method allows us to track both translated BicDR-A and BicDR-B. Through sequencing, we confirmed that the BicDR ORF was fused seamlessly with the eGFP ORF. The successful ORF fusion and the expression of the predicted fusion protein were also confirmed by Western blotting of embryonic extracts which revealed the GFP expression as part of a 120 kDa polypeptide (Figure 4A).

Immunolocalization of BicDR::GFP in embryos revealed that the apical side of salivary gland cells stained very strongly from stage 13 on (white arrow in Figure 4B, higher magnification shown underneath). Additionally, individual cells displayed staining signals in a metameric manner along the lateral side of the embryo. These signals were most intense during stages 11-14 (Figure 4C, top). By co-immunostaining of BicDR::GFP and the nuclear neuroblast marker Asense (Ase), which is expressed in all sensory organ precursor (SOP) cells and their progeny^41^, the BicDR::GFP expressing cells appear to concentrate in the region of the Ase positive cells but do not clearly overlap with them (Figure 4C).

In preparation for metamorphosis, this first set of sensilla in the embryos degenerates during the late third larval instar^44^. Because the adult sensory organs, the bristles, are built similarly and give rise to larger structures, we tested if BicDR::GFP is also expressed in the second set of SOPs, which are produced by the imaginal wing disc histoblasts that form the adult epidermis^44^, we repeated the co-immunostaining with wing discs of third instar larvae that express the endogenous BicDR as a GFP fusion. Although we could not observe an increase of the signal specifically in the SOPs, we could confirm that the BicDR::GFP signal is present in the dorsal posterior area of the wing discs that will form the notum with the SCps and SCas of the adult fly (Figure 5).

**Figure 5.**
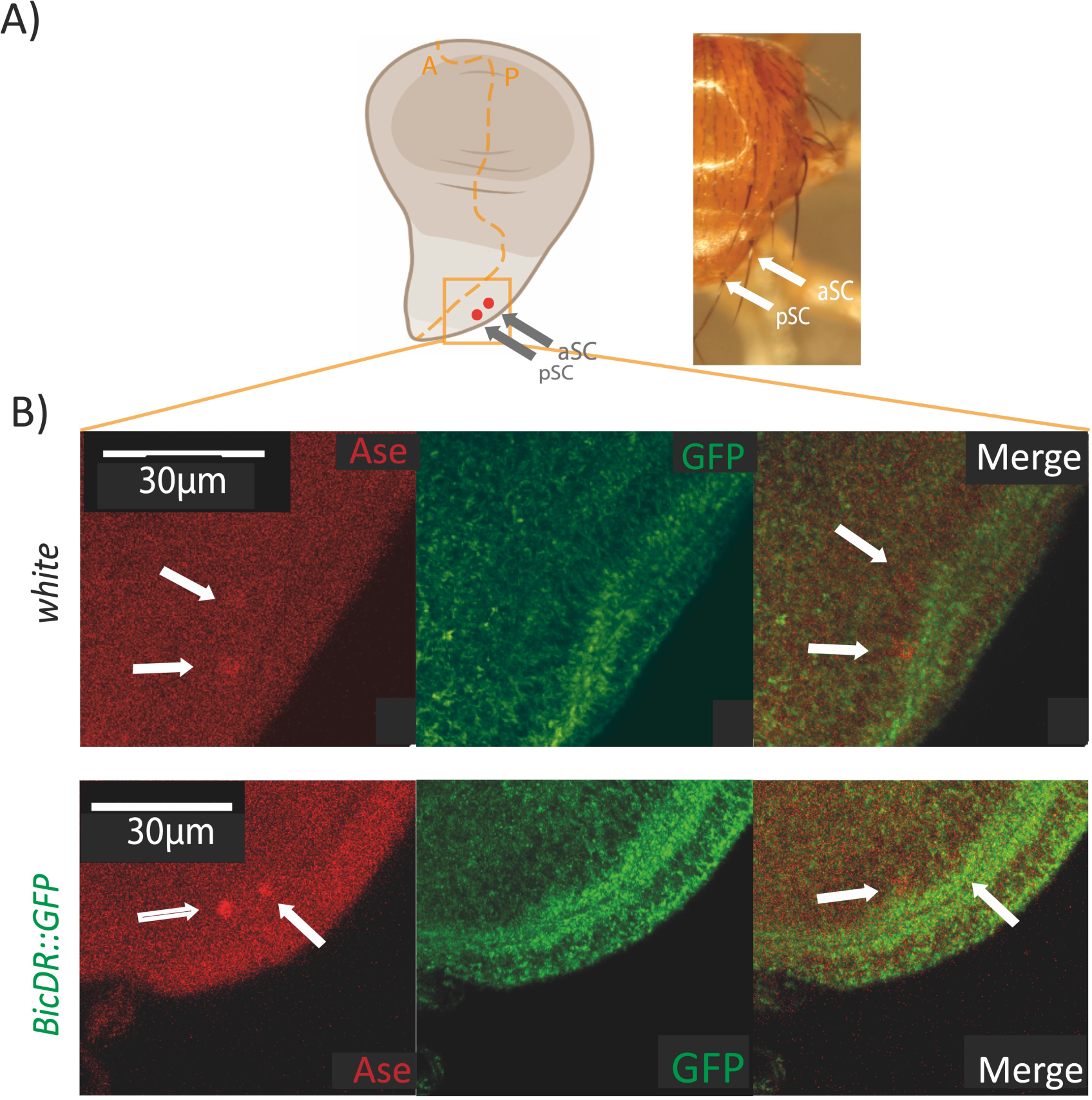
The expression of BicDR::GFP in dorsal wing discs of third instar larvae. **A)** L3 wing disc showing the position of cells giving rise to dorsal macrochaetae of the adult. **B)** BicDR::GFP staining signal in the control (white) and in larval discs expressing BicDR::GFP. The staining for GFP is above the background in the posterior part of the wing discs, but it does not specifically accumulates in the SOPs that will form SCa or SCp.

### *BicD* and *BicDR* contribute to localizing Rab6 to the tip of the mechanosensory bristles

To further understand bristle development and the impact of *BicD* and *BicDR* on it, pupal dorsal tissue containing the developing bristles was dissected 40 to 44 hours after pupation, fixed, and stained. In this way, hemizygous *BicD^PA66^* samples with only one functional copy of *BicDR* were compared to *BicD^PA66^* animals with two functional *BicDR* copies and to controls that were wild-type for *BicD* and *BicDR*. Investigating the F-actin structure of the samples allowed us to compare the morphology and length of the macrochaetae (Figure 6). Comparing the pupal bristle length in the mutants with the wild type showed that the mutant bristles were somewhat shorter but that there were no significant length differences (Figure 6A). This indicates that the short bristle phenotype observed in *BicD*; *BicDR* double mutant flies evolved at a later stage of development. Although similar in length, the actin cytoskeleton of the double mutant scutellar bristles displayed abnormalities. 6 out of 7 mutant scutellar macrochaetae showed an irregular arrangement of the actin bundles, and obvious gaps could be observed (Figure 6B). This phenotype could not be observed in the control pupae (0 out of 4 scutellar macrochaetae) nor in hemizygous *BicD^PA66^* animals. While such gaps in the actin bundles are reminiscent of the chitinization process of the bristles, this does not appear to to be the reason because all pupae were only 40 to 44 hours into pupation. The breakdown of the bundles by chitinization, however, begins only 48 hours after pupation and it shows initially narrow longitudinal gaps between modules and these become wider as the bristle ages. Only in 53 hours old pupae such breakdown becomes clearly recognizable^45^.

**Figure 6.**
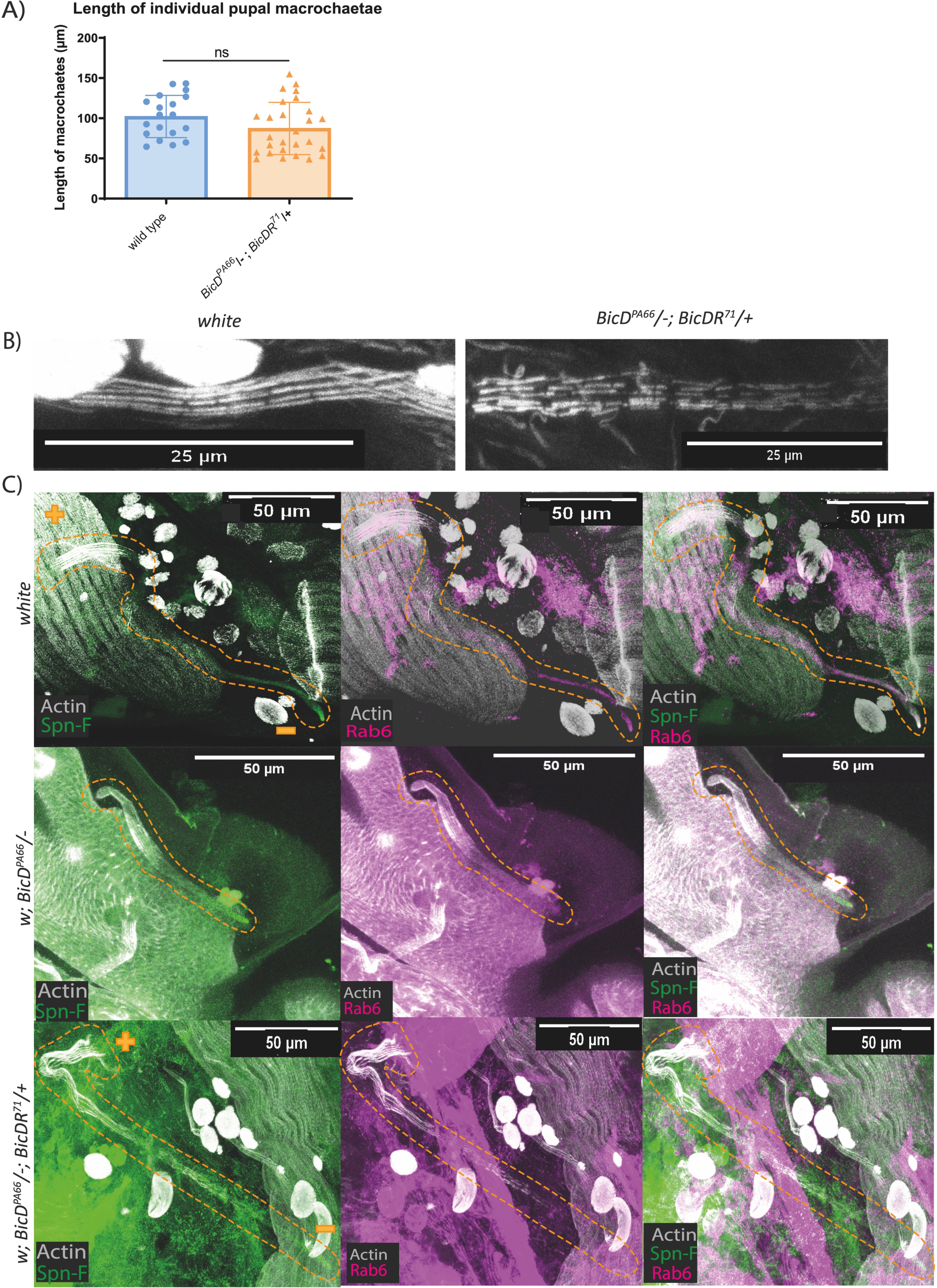
*BicD; BicDR* mutants display irregularly lined up actin modules in scutellar macrochaetae. **(A)** The length of single pupal macrochaetae was measured in the *white* control and the *BicD^PA66^*/−; *BicDR\**/+ double mutants. No significant dissimilarities between the two groups were found at this stage of bristle development. **(B)** Stained scutellar macrochaetae in controls and double mutants (grey: F-Actin). **(C)** *BicD* and *BicDR* are needed to localize Rab6 at bristle tips in wild-type scutellar macrochaetae. This accumulation is impaired in *BicD*, *BicDR* double mutants (grey: F-Actin, green: Spn-F, pink: Rab6). The genotype of the sample is listed on the left side. All macrochaetae originate in the upper left corner (indicated with a “+”) and grow downwards to the lower right corner (indicated with a “-”). The tip is visualized with the staining of Spn-F.

Rab6 is known to be a Notch modifier that influences the development of the mechanosensory bristles on the head, notum, and scutellum. The *Rab6* phenotype also results in aberrant bristle length and bristle tips that have very mild and disorganized ruffling^46^. This phenotype resembles the short bristle phenotype observed in *BicD*; *BicDR* double mutants. Additionally, Schlager and colleagues described a physical interaction between Rab6 and BicDR in *Danio rerio*^47^, and Januschke et al. an interaction between *Drosophila* Rab6 and BicD^15^. We, therefore, examined the Rab6 distribution in the macrochaetae of *BicD^PA66^*/− and *BicD^PA66^*/−; *BicDR\**/+ mutants. For this, we stained the pupal dorsal tissue for Rab6. As seen in Figure 6C, the Rab6 signal is for the most part present at the tip of wild-type scutellar bristles, similar to the microtubule minus-end-marker Spn-F^2^. In *BicD^PA66^*/− bristles, the Rab6 signal does either not appear, just like in *BicD*; *BicDR* mutants, or it is visible, but evenly distributed throughout the bristle shaft without a discernable distal tip accumulation (Figure 6C). Our data suggest that transport aberrations to the distal tip in combination with a defective actin cytoskeleton prevent normal bristle construction in *BicD^PA66^*/−; *BicDR\**/+ mutants and that the localization of Rab6 at the bristle tip is driven by both *BicD* and *BicDR*.

### *BicDR* and *BicD^PA66^* in localizing Spn-F to bristle tips

Spindle-F (Spn-F) is a microtubule minus-end marker that affects oocyte patterning and bristle morphology in *Drosophila*^48^. *Spn-F* mutants eclose with shorter and thicker bristles. Scanning electron micrographs of the bristles revealed that the mutant bristles have branching tips and that the direction of elongation is sometimes perturbed^48^. Spn-F functions at the distal tip of the growing bristle and is involved in the regulation of the shuttling movement of recycling endosomes and cytoskeletal organization^49^. We analyzed the potential requirement for *BicD* and *BicDR* for the localization of Spn-F to and within the shaft of the bristle cells (Figure 6C, Figure 7). The normal asymmetric localization to the tip of the macrochaetae allowed us to assess the contribution of *BicD* and *BicDR* to this microtubule minus-end transport process. One measure for the establishment of the polarity of the bristles is the “tip index”: a line scan from the bristle shaft to the distal tip establishes a plot profile from which the maximum intensity along the bristle length is determined. The “tip index” is defined as the relative position of the pixels that exceed 50% intensity along the bristle axis^49^. This index is used to quantify the asymmetric localization of a protein within the bristle cell. If a signal is completely localized at the bristle tip, the tip index will have a value of 100. If the signal remains in the cell body and stays absent from the bristle, the value of the tip index is 0^50^. This measurement confirmed that the Spn-F signal is significantly more concentrated at the tip of the macrochaetae of control pupae, while this signal tends to appear diffusely throughout the whole cell in *BicD^PA66^*/− and w; *BicD^PA66^*/−; *BicDR^71^* / + bristles (Figure 7A and 7B). The tip index in control macrochaetae had a value of 33, while the value in w; *BicD^PA66^*/−; *BicDR^71^*/+ bristles was 18. Similar to this, the tip index in w; *BicD^PA66^*/− bristles was 16. These results suggest that *BicD* is necessary for the localization of Spn-F to the distal tip and that its function is not redundant with *BicDR*.

**Figure 7.**
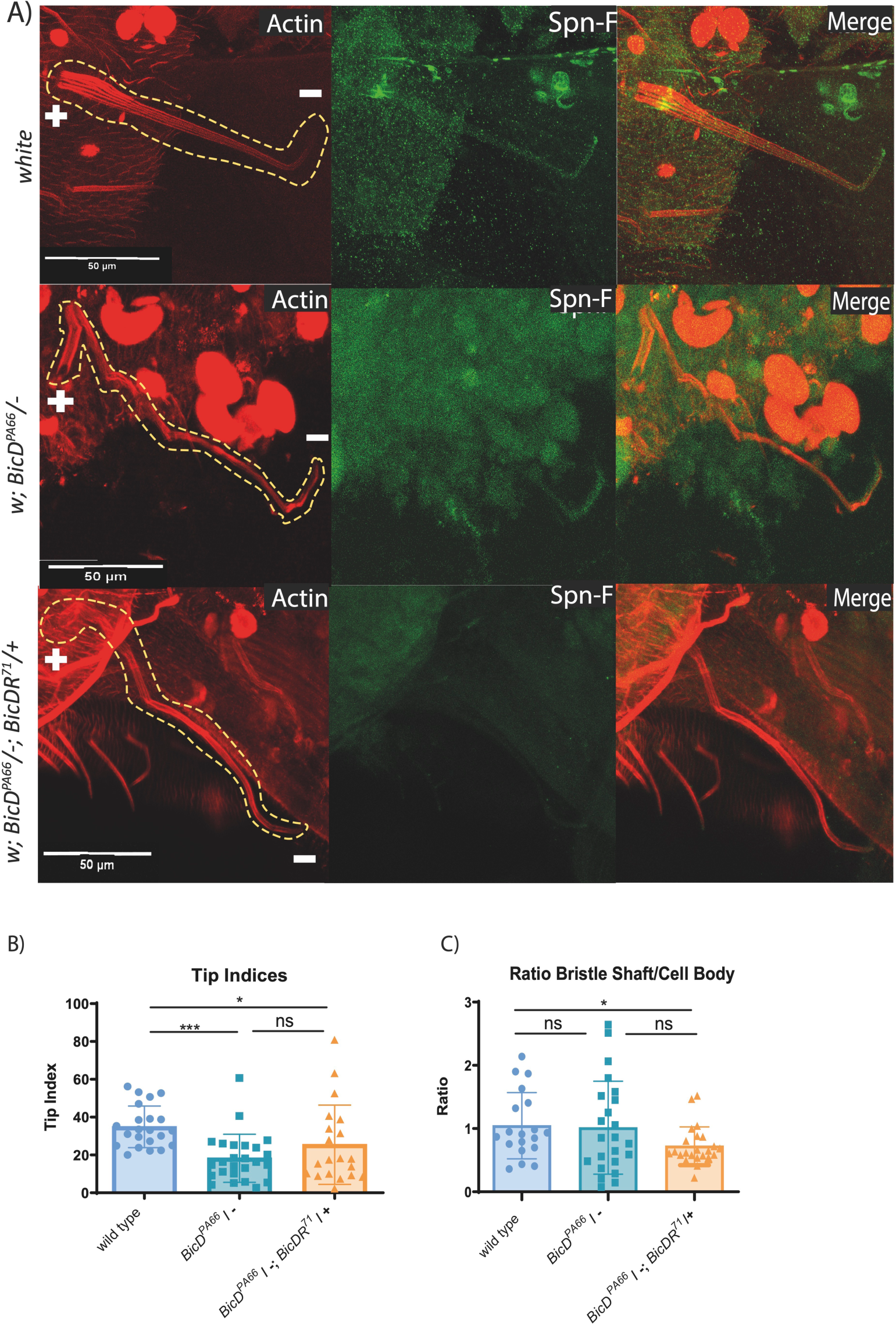
Pupal bristles of *BicD*; *BicDR* double mutants show impaired Spn-F localization at the bristle tip. **(A)** Stained macrochaetae (red: F-Actin, green: Spn-F). The genotypes of the samples are listed on the left side. All macrochaetae originate in the upper left corner and have their tips pointing downwards. The localization of Spn-F at the bristle tip is much weaker in *w*; *BicD^PA66^*/*Df7068*; TM6B/+ and *w*; *BicD^PA66^*/*Df7068*; *BicDR^71^*/+ pupae. This observation could be confirmed with the calculation of the tip index shown in **(B)**. The average tip index of the mutants is significantly lower than those of the control group. **(C)** The ratio of Spn-F signal in the elongated bristle shaft versus its cell body is significantly lower in *BicD^PA66^*/−; *BicDR\**/+ animals.

Complementary to the previous signal distribution is that the Spn-F signal ratio in the bristle shaft/cell body changes between *BicD* and *BicD*; *BicDR* double mutants (Figure 7C). To quantify this, we measured the average signal strength of an area on a plane in the center of the bristle shaft and divided this by the average signal strength of an area of the same size drawn on a plane through the bristle cell body, directly under the bristle root. While w; *BicD^PA66^*/− bristles show a wider distribution of this ratio, the ratio decreases significantly in w; *BicD^PA66^*/−; *BicDR^71^*/+ bristles in comparison to the wild type. One might, for instance, expect to find such a distribution if BicDR is more involved in localizing Spn-F to the periphery of the macrochaetal cell body and BicD more for transporting it along the bristle shaft towards the bristle tip.

At the developmental stage when we observed these localization differences, the length of the mutant macrochaetae was not significantly reduced yet (Figure 6A). It thus appears that the lack of proper localization of Rab6 and Spn-F affects bristle construction at the subsequent stage, possibly during the chitinization and the associated deconstruction of the actin filament.

### EF1γ is found in BicDR complexes and *EF1γ* enhances the bristle phenotypes of *BicD* and *BicdR*

To learn more about the mechanisms through which BicDR contributes to transport processes in general, we used the C-terminally tagged endogenous *BicDR* (*BicDR::GFP*) and performed immunoprecipitations with an anti-GFP antibody using extracts from 10-16 hours old embryos. We also mutated the endogenous gene into a *BicDR^K555A^::GFP* gene and used it as a control because it might allow us to distinguish between the cargo that binds through the K555 region and other interacting partners of BicDR. A *white^-^*strain with an untagged *BicDR^+^* was used as a negative control. The search for interacting proteins was performed in 2 different ways. First, in triplicate experiments, embryos were lysed and immunoprecipitated with anti-GFP antibodies. A proteomic analysis was then performed directly on the precipitated fractions. Second, embryos were lysed in duplicates, immunoprecipitated as before, and the resulting proteins were separated by SDS-PAGE. The gel was stained by Coomassie blue and only those bands found in the tagged *BicDR::GFP* fraction and not in the *BicDR^K555A^::GFP* samples were excised and analyzed. Gel slices from the corresponding position of the control samples were also analyzed.

The results of the first IP experiment defined 25 potential BicDR::GFP interactors with a p-value ≤0.05 and log_2_FC ≥1.0 (Supplementary data file S1). Of these, 7 were also found in BicDR^K555A^::GFP samples, indicating that these are binding partners that depend less on K555. Out of the remaining 18 candidates, different bristle phenotypes had already been described for 4 mutants (*tou*, *RpS17*, *RpL27A,* and *RpL12*)^51–53^ while RNA binding activity had been observed for RpS5b^54^ (Tables S1). Whereas ribosomal proteins are a common contaminant in IPs, mutations in ribosomal protein genes lead to impaired bristle development and show a haploinsufficiency phenotype that is seen as evidence for a very high protein synthesis required for bristle development^55^. It is therefore also possible that BicDR interacts with ribosomes. *Tou*, on the other hand, is a transcription factor that activates proneural gene expression^53^ and has also been found in a gain-of-function screen for genes that affect external sensory organs^56^. The overexpression of different *tou* alleles results in excess scutellar and dorsocentral macrochaetae^57,58^.

Other noteworthy candidates identified in this IP are Rac1 and *morpheyus* (*mey*). Although identified with only a few counts, Rac1 is significantly enriched in the IP with the wild-type BicDR::GFP peptide. It is associated with axial outgrowth^59,60^ and highly expressed in the salivary glands^61^, where it functions to control lumen size. By the fact that it controls epithelial tube length and formation through apical secretion and regulation of cytoskeletal dynamics in tracheae and embryonic salivary glands^62,63^, the functions of Rac1 are especially interesting for further analysis of *BicDR* expression and function. Furthermore, bristle defects are also caused by overexpressing mutant *Rac1* alleles^64^. *mey* is expressed in the same tissues as BicDR, namely the tracheal system and salivary glands. Although no *mey* phenotype has been described yet in these tissues, it has been reported that knock-down mutants develop abnormal touch responses during the larval stage. This could point to a defect in larval mechanosensory organs.

In the second approach with gel-purified bands, the larger sample size yielded 179 interacting proteins in the tagged wild-type BicDR::GFP IP that were not present in the IP of the tagged mutant (Figure 8A; Supplementary data file S2). Because the *BicDR^71^* chromosome showed a bristle duplication indicative of a problem in Notch-dependent binary cell fate acquisition^65^, and because Notch signaling also depends heavily on cytoplasmic transport, we searched among the proteins identified in the BicDR::GFP IP for known trafficking regulators of the Notch receptor (Table S2). Origin recognition complex subunit 6, Vacuolar H^+^ ATPase subunit 68-2, Vacuolar H^+^ ATPase 26kD E subunit, Rumi, Par-6, and Ef1γ are all Notch-trafficking regulators that were absent in the control IPs but detected in the BicDR::GFP IPs. Except for *par-6*, a loss-of-function mutation of all the genes for these candidate interactors results in bristle-loss^65^.

**Figure 8.**
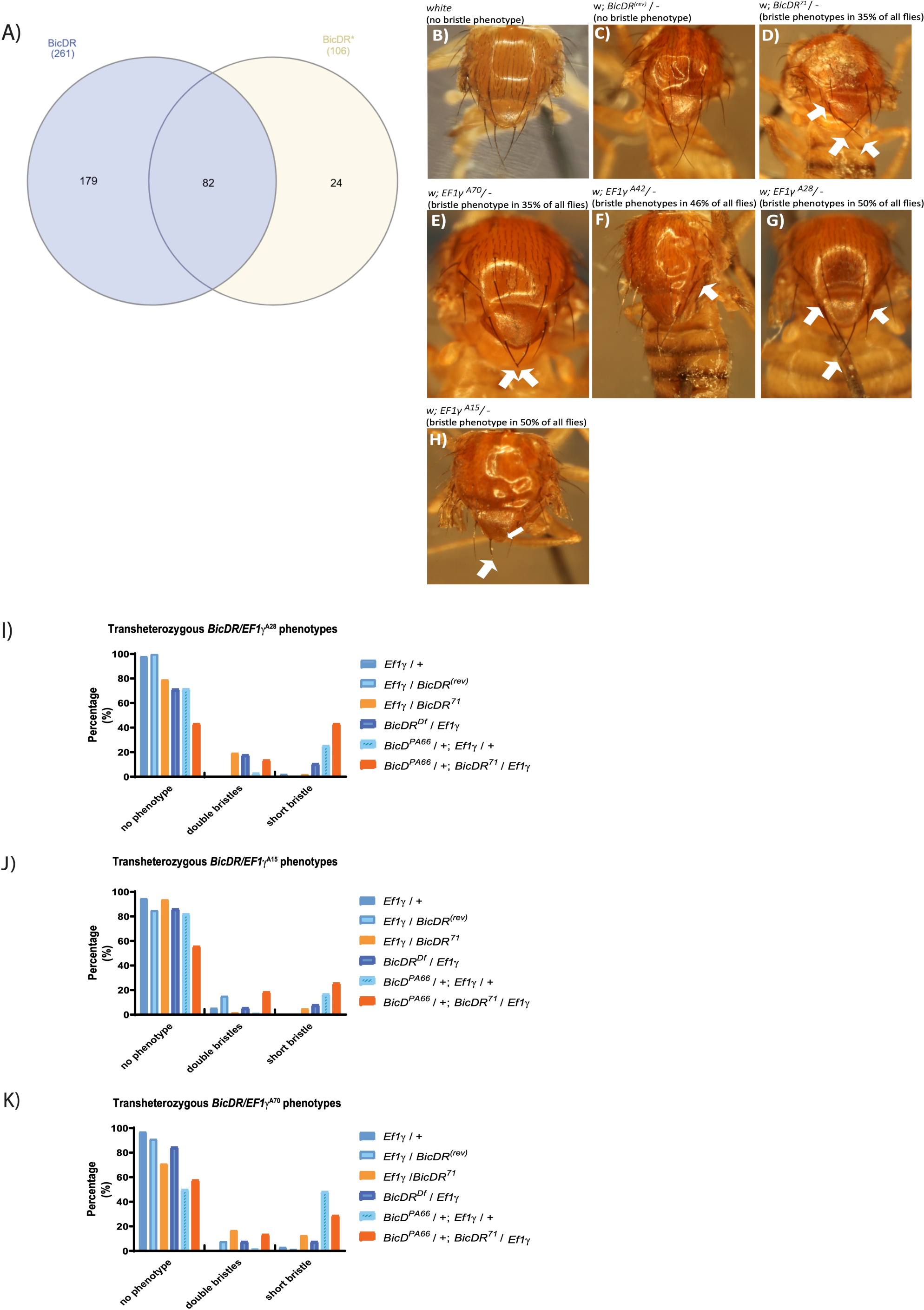
(A) Proteins identified in the cut-out gel bands from the tagged BicDR and BicDR^K555A^ immunoprecipitations. **(A)** A total of 285 potential binding partners were identified. 82 proteins were found in both samples, while 179 proteins were found only in the wild-type BicDR::GFP IP. **(B)-(H)** Resemblance of phenotypes compared to the **(B)** *white* control (0% observed bristle phenotype), **(C)** *BicDR^(rev)^*/− control (0% observed bristle phenotype), **(D**) *BicDR^71^*/− (35% of adult flies display such bristle phenotypes), **(E)** *EF1γ^A70^*/− (35% of flies display shorter bristles), **(F)** *EF1γ^A42^*/− (46% of adult flies ecloses with additional bristles), **(G)** *EF1γ^A28^*/− (50% of adult flies have additional and shorter bristles), **(H)** *EF1γ^A15^*/−(50% of all flies have shorter bristles). Note that the *white* control and 93% of the *BicDR revertants*, *BicDR^(rev)^*/*-,* eclosed without a visible bristle phenotype. A total of 7% of *BicDR^(rev)^*/− animals contained an additional aSC bristle. Flies with the genotypes *BicDR^71^*/−, *EF1γ^A70^*/− and *EF1γ^A51^*/− showed shorter pSC macrochaetae. Additionally, 21% of *BicDR^71^*/− animals eclosed with an extra aSC bristle, a similar frequency as observed with the alleles *EF1γ^A42^*/− and *EF1γ^A28^*/−. (**I-K)** Effect of combining heterozygous *EF1γ^A70^*, *EF1γ^A42^* or *EF1γ^A28^* with heterozygous *BicD* and *BicDR* alleles shows strong genetic interactions between heterozygous *BicD*, *BicDR,* and *EF1y* alleles. Frequency of mutant phenotypes observed in double and triple heterozygous combinations. Different mutant combinations containing a *BicDR*, BicD^PA66^*, and *EF1γ\** allele eclosed with different bristle phenotypes. The frequency of animals that eclosed with a short-bristle phenotype is significantly higher if the animals carry a *BicD^PA66^* and *BicDR^71^* allele except for the combination with *EF1γ^A70^* where the frequency of flies with short bristles was the highest in *BicD^PA66^*/+; *EF1γ^A70^*/+ (30%). Noticeable is that 19% of the flies with the genotype *BicD^PA66^*/+; *BicDR^71^*/*EF1γ^A28^* eclosed with short bristles, even though the allele *EF1γ^A28^* /− induces additional bristles.

The translational regulator Ef1γ appeared also interesting because its mutants display a bristle phenotype and it was immunoprecipitated at the highest amount among the identified potential binding partners of BicDR. Aside from its function in translation, EF1γ is known to negatively regulate the transport of several classes of membrane organelles along microtubules^27^ and for its interaction with keratin bundles in mouse fibroblasts^66^. For these reasons, we further investigated the interaction with EF1γ. To test whether the two genes might act in the same pathway, we first compared their mutant phenotypes (Figure 8B-8H). While the alleles *EF1γ^A42^* and *EF1γ^A28^* induced additional aSC macrochaetae at either only one or both sides of the notum (Figure 8F and 8G), the mutants *EF1γ^A70^* and *EF1γ^A15^* eclosed with stubble-like pSC macrochaetae, similarly to hemizygous *BicDR^71^* flies (Figure 8E and 8H and Figure 8D).

Flies transheterozygous for *BicDR^71^* and *EF1γ* are viable and 2-13% of them displayed shorter pSC or aSC macrochaetae (*BicDR^71^* and *EF1γ^A28^*: 2%; *BicDR^71^*/ *EF1γ^A15^*: 5%; *BicDR^71^/ EF1γ^A70^*: 13%; Figure 8I, 8J and 8K). This effect could not be observed in *BicDR^(rev)^* / *EF1γ^A28^* mutants or heterozygous *EF1γ^A28^* animals. The phenotype was significantly more prominent if the flies were transheterozygous for *BicD^PA66^* and *EF1γ^A28^*: 23% of the animals showed at least one shorter bristle. This went up to 48% with *BicD^PA66^* / +; *EF1γ^A70^* /+, whereas in *BicD^PA66^* / +; *EF1γ^A15^* /+ flies 17%; showed at least one short bristle (Figure 8I-K). To test if a mutant *BicDR* allele enhances the phenotype of transheterozygous *BicD^PA66^*; *EF1γ^A28^* mutants even further, we generated flies that were heterozygous for all three genes. 44% of all *BicD^PA66^*/+; *BicDR^71^/ EF1γ^A28^* eclosed with at least one shorter bristle (26% in *BicD^PA66^*/+; *BicDR^71^/ EF1γ^A15^* animals and 30% in *BicD^PA66^*/+; *BicDR^71^/ EF1γ^A70^*).

In summary, we conclude that except for allele *EF1γ^A70^*, the proportion of animals with shorter bristles is significantly higher if they are heterozygous for all three mutants, *BicD^PA66^*, *BicDR^71^*, and *EF1γ*, indicating that all three genes are functioning in the same direction and contribute to proper macrochaetae development. This appears surprising because BicD and BicDR help to perform MT transport whereas *EF1γ* negatively regulates it. The observed type of genetic interaction can be explained if *EF1γ* performs its function at the bristle tip and negatively regulates organelle transport there, allowing the organelles to perform their function at the tip. Unfortunately, the antibody localization of EF1γ did not allow us to test the distribution of EF1γ in the pupal bristles. Presumably because of the high signal levels in all tissues, one would need to use a more complex approach to test this.

Also in this situation, the female flies were much more affected by this phenotype than the males and additional bristles could be observed at low frequency in the mutants *BicD^PA66^*/+; *BicDR^71^/ EF1γ*, but also in the controls (*EF1γ*/+ and *EF1γ* / *BicDR^(rev)^*). The similar bristle phenotype, the genetic interaction between *EF1γ* and *BicDR*, and the fact that BicDR::GFP and EF1γ co-precipitated posed the question of whether they directly interact. However, a yeast two-hybrid assay did not detect a direct interaction between BicD, BicDR, or EF1γ (Figure S2). This suggests that the interaction between BicDR and EF1γ is mediated by another complex component. Additional interactors from the same screen are Arp2 and Arp3 (Table 2). The Arp2/Arp3 complex is involved in the organization of the actin filaments, a structure that is affected by the reduced *BicD*; *BicDR* function (Figure 6). Because EF1γ was the top hit in this group and the genetic interaction assay testing for combined haploinsufficiency showed a stronger interaction with the *EF1γ* alleles (Figure 8), we, however, decided to focus on *EF1γ* for the proof of principle in the present study.

## Discussion

We found that *BicDR* is not an essential gene, but its modest phenotypes in the trachea (Figure 4) and bristles (Figure 3) reveal important functions, particularly for life outside the laboratory setting. Additionally, one functional *BicDR* copy is essential for viability in a hypomorphic *BicD* background. In these animals with reduced *BicD* activity and only one functional copy of *BicDR,* the remaining combined activities of *BicD* and *BicDR* activity are not sufficient to develop bristles properly. Here we showed that the reduced activity of *BicD* and *BicDR* affects the Rab6 and Spn-F localization in the growing bristle, linking BicD and BicDR to the dynein-dependent microtubule transport of vesicles and bristle factors to their proper position in the bristle where they perform their function.

A different defect in the development of the bristle, the formation of a twin bristle on the notum, was seen in 21% of hemizygous *BicDR^71^* flies (Figure 2D). This was mostly an additional aSc bristle with a hair and socket of its own. This hinted at a failed cell fate acquisition after the division of pI cells that can result from gain-of-Notch signaling in the cell divisions leading to the sensory organ^67^. This connection was also attractive because the Notch trafficking regulator EF1γ^67^ was a top hit for BicDR interacting proteins and transheterozygous EF1γ/BicDR^Df^ also showed bristle duplications. *BicDR^Df^* lacks, aside from BicDR, 8 other genes. However, the evaluation of the cause of this phenotype became too challenging for now because animals in the control group *BicDR^(rev)^*/−, a wild-type revertant generated by hopping out the P-element insertion that caused the *BicDR^71^* allele after an imprecise excision, also showed twin bristles in 7% of the animals. On the other hand, excision mutants and revertants that were generated with the P-element that had inserted in the 5’ region of *BicDR* (Figure S1) did not show this phenotype. A possible interpretation might be that the P-element chromosome had acquired a second hit that supports bristle duplications. Such second-site hits are common and often caused by local transpositions, which would explain why the *BicDR^Df^* chromosome also showed this interaction with EF1γ (Figure 8I-K). Because of the difficulty of resolving this issue, we decided to focus for now mainly on the bristle growth phenotype to study the function of *BicDR*.

The second coiled coil domain of BicD ensures that the adaptor protein remains inactive if no cargo is bound. For this, the cargo-binding third coiled coil domain folds back onto the second coil coil, thereby blocking the dynein interaction site^68^. This mechanism ensures that the BicD-transport machinery does not run unloaded along microtubules. BicDR lacks this second coiled coil domain, suggesting that BicDR itself does not have an activated or inactivated state or controls this through a different mechanism. A second dissimilarity between *BicD* and *BicDR* is the big difference between their expression levels. According to Flybase^69^, *BicDR* is mainly expressed in tracheae, gut, salivary glands, and carcass tissue, while the expression in other tissues remains at low levels. Although there is some overlap with the expression of *BicD*, the expression of *BicDR* was described to remain low in the tissues where the consequences of *BicD* mutations have been described. Such as in the ovary, the young embryo, and the nervous system^70–72^. This allows the assumption that the expression of BicDR at low levels does not necessarily contribute to cargo transport the same way as BicD does. How could BicDR then support BicD? It does not appear to dimerize with a BicD subunit based on immunopurification or yeast 2-hybrid results. *BicD* is more important in large cells where it transports cargo over very long distances. *BicDR* might be specialized for moving cargo for local transport over short distances (e.g., in the tracheal cells). In cells where both are expressed, BicDR may then make the cargo more accessible for long-distance transport by BicD. This seems consistent with the function we found in the growing bristle shaft, where BicDR seems more involved in bringing Rab6 and Spn-F from the cell body to the shaft and BicD to transport them toward the tip (Figures 6-7). While the proper localization of Spn-F depends on *BicD* and *BicDR*, we found that in the sensitized background (*BicD^PA66^*/−) both copies of *BicDR* are needed to move normal levels of Spn-F from the cell body into the bristle shaft (Figure 7C). On the other hand, full *BicD* activity is not required for this step (Figure 7C) but is required to obtain normal levels of Spn-F at the bristle tip (Figure 7B).

*spn-F* is needed for the localization of Hook at the bristle tip^2^ and *hook* is not only required for endocytic trafficking within the eye and the nervous system but also at the bristle tip. Since there is evidence that endocytosis is responsible for the polarized transfer of lipids and membrane proteins, which again is necessary for the polarization of the bristle cell^73^, our results point to an important contribution of BicD and BicDR to bristle development by localizing Spn-F to the tip. With Spn-F also being part of the IKKε-jvl complex, which regulates the shuttling movement of recycling endosomes and cytoskeletal organization^50^, the lack of Spn-F in this complex would interfere with the shuttling regulation of motor proteins at the molecular signaling centers^50^. This also prevents the transport of Rab-positive vesicles. Mutations in Rab6 and Rab11, members of the Rab protein family that mediate endocytic intracellular vesicle trafficking, lead to impaired bristle growth^9,74,75^. The localization of the Rab6 signal to the bristle tip is in line with the description of the distal tip being the signaling center for bristle elongation and thereby the most dynamic part of the polarized cell^49^. The failure to localize Rab6 at the distal tip in hemizygous *BicD* hypomorphs with either one or two functional copies of *BicDR* indicates that this active exo- and endocytosis at the bristle tip is impaired. This suggests that the developing bristle requires *BicD* and *BicDR* for the bristle construction by localizing parts of the endocytic compartments to the bristle tip.

Rab11 contributes to the construction of the bristle by inserting chitin synthase into the plasma membrane and thereby allowing bristle chitinization^77^. The complete lack of chitin synthase in *Rab11^-^*bristles, the bristles not only appear shorter but collapse completely.The fact that we did not observe clear differences in bristle length during the pupal stage of *BicD*; *BicDR* double mutants also suggests that the limiting step in these animals is the construction of the final macrochaetae with their complete chitinization. Because the Rab11 phenotype is quite different from the *BicD*; *BicDR* double mutant bristle phenotype, we initially did not focus on Rab11. However, with the knowledge gained from this study, we might be able to explain the *BicD*; *BicDR* double mutant bristle phenotype also by a requirement for proper Rab11 localization in the bristle shaft and tip. Reduced transport of Rab11 or a transport factor toward the tip might cause a polar reduction of chitin incorporation towards the tip, causing an increased probability of breakage in the distal parts under reduced *BicD* and *BicDR* activity.

Actin modules in the bristle shaft are central to the construction of the bristle. *BicD* and *BicDR* promote the accumulation of Rab6-vesicles (and possibly other vesicles) at the bristle tip and contribute to the proper build-up of the actin modules. The disorganized F-actin network seen in the mutants can either be caused by insufficient maintenance and stability of the F-actin, resulting in premature breakdown of the modules or incorrect alignment of already formed bundles. These defects can be expected to prevent normal chitinization. Because shorter macrochaetae on the head or notum of the pupae did not show this phenotype, we assume that the impaired transport of bristle components or organizers over the long distances to the tip of the long scutellar macrochaetae cannot keep up with the turn-over rate of actin filaments or provide the needed stabilization factors, causing degeneration of their cytoskeleton, breakage of the bristle shaft, and shorter or thinner bristles^45,76^.

Our results demonstrate how directed transport contributes to the organization of asymmetric cells. The microtubule transport, which localizes factors that organize cellular functions, connects directly and indirectly to vesicle trafficking and the stability of the actin cytoskeleton. We could show that *BicD* and *BicDR* contribute together to this directed transport and the development of the bristles in a partially redundant manner. Our results led to the hypothesis that BicD might be more specialized for long-haul transport and BicDR more for short-distance, local transport. Future studies should test this hypothesis.

## Acknowledgments

We thank S. Bullock, L. Rabinow, A. K. Satoh, U. Abdu, S. Luschnig, the Bloomington *Drosophila* Stock Center (NIH P400D018537), and the Developmental Studies Hybridoma Bank (created by the NICHD of the NIH and maintained at The University of Iowa, Department of Biology, Iowa City, IA 52242) for constructs, fly stocks and antibodies. We thank FlyBase (U41HG000739) for the *Drosophila* genomic resources. Special thanks go to D. Beuchle and C. Elci for their excellent technical help.

This work was supported by funds from the Swiss National Science Foundation (SNF, project grants 31003A_173188 and 310030_205075; www.snf.ch) and the University of Bern (www.unibe.ch) to BS.

## Legends for supplementary Figures

**Figure S1.**
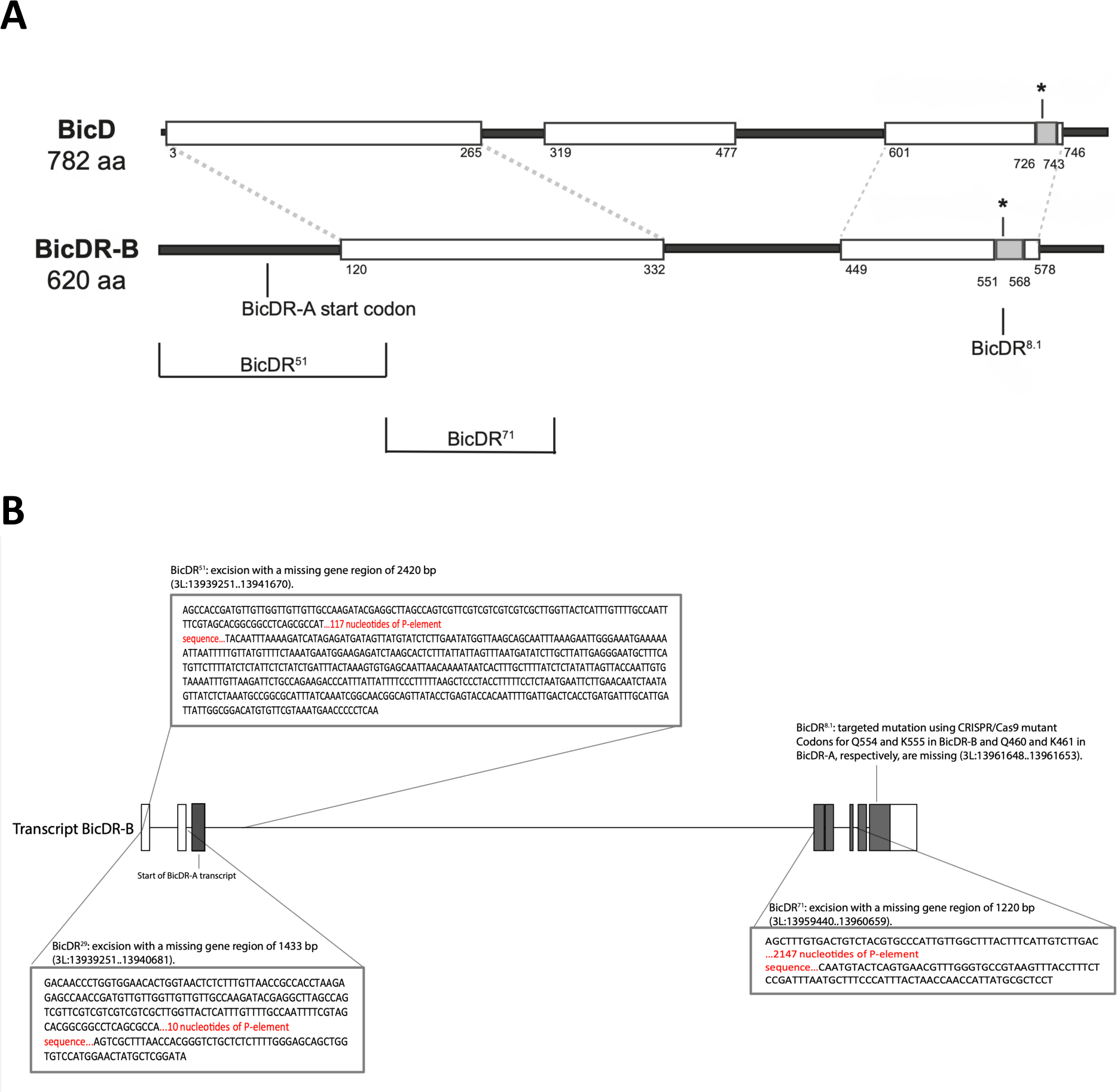
Comparing BicD and BicDR structures and the *BicDR* gene. **A)** Structure comparison of the *Drosophila* proteins BicD and BicDR. Shown in open boxes are the coiled coil domains, in filled boxes the sequence with the highest homology. The lysine with a key role in cargo attachment is indicated with a star. It is localized at position 730 in BicD, at position 555 in BicDR-B, and 461 in BicDR-A. The regions altered in the different alleles are indicated. The hemizygous deletion *BicDR^8.1^* removes precisely the two codons Q554 and K555 in BicDR. **B)** Structure of *Drosophila BicDR-A* and *–B* mRNAs and the excision mutants. The gray boxes frame the parts of the gene that have been removed by the imprecise P-element excision. The sequences in the box show in detail where the excision breakpoints are. Sequences in red are P-element leftovers. The excision *BicDR^29^* misses the 5’ UTR region, without impairing the first protein-coding exon, while *BicDR^51^* misses the first exon of *BicDR-A* and *–B* and the 5’ UTR region as well. *BicDR^71^* is the only excision that removes exons 2, 3, and 4 but leaves exons 5 and 6 intact. *BicDR^8.1^* is a CRISPR mutant with deleted Q554 and K555.

**Figure S2. *Drosophila* BicDR does not interact with EF1γ, Egl, or BicD (A)** The activation domain is indicated on the left and the binding domains are on the right side. The latter is drawn where these activation domains were plated out. (-LW): grown on medium selecting for the two plasmids, (-LWH): selective plates on which cells with the activator domain and the DNA binding domain clones can grow if they weakly interact. (-LWH) with 20 mM 3-aminotriazole [3-AT]: selective plates on which cells with the activator domain and the DNA binding domain clones can grow if they strongly interact. FL: full length, CTD: C-terminal domain, BD: binding domain, AD: activation domain. BicD and Egl were used as positive controls since their interaction had already been demonstrated^79^. Here, the full-length protein of BicDR, fused to the activation domain, binds to the C-terminal domain of BicDR as well as to the C-terminal domain of BicD. However, this result is not confirmed under stringent conditions, since neither the full-length BicD protein nor the C-terminal domain of BicD seems to bind to BicDR in immunoprecipitations nor according to the yeast 2-hybrid assay. Thus, these results seem to indicate that the direct interaction between BicDR FL (AD) and BicD (BD) resulted from the nonspecific entanglement of the coiled coil domains^80^.

**Table S1.** Proteins that were significantly enriched in the tagged BicDR::GFP Ips in comparison to the wild-type negative control IPs. Genes encoding these proteins are listed according to their level of significance of enrichment over the control. The abbreviations used are “control” for the *white* control, “BicDR” (BicDR::GFP), and “BicDR^K>A”^ (BicDR^K555A^::GFP). Hits enriched in tagged wild-type BicDR compared to BicDR^K>A^ are listed as potential cargo, whereas peptides enriched in tagged wild-type BicDR and BicDR^K555A^ IPs are listed as potential non-cargo interactors. GN is the abbreviation for gene name. The iBAQ equals the sum of all peptide intensities divided by the number of observable peptides of a protein^1^. log^2^FC is the logarithm of the mean ratio between the two groups and the adjusted p value (adj. pVal) highlights the factor level comparisons within a family that are significantly different^2,3^. −1 and −2: indicate different biological replicates.

| GN | Sequence coverage [%] | iBAQ | log2FC BicDR - control | log2FC BicDR<sup>K>A</sup> - control | log2FC BicDR<sup>K>A</sup> - BicDR | Adj. pVal BicDR - control | Adj. pVal BicDR<sup>K>A</sup> - control | Adj. pVal BicDR<sup>K>A</sup> - BicDR | Pot. Cargo | Pot. Non-cargo interactor |
| --- | --- | --- | --- | --- | --- | --- | --- | --- | --- | --- |
| BicDR | 84.7 | 1.70E+09 | 11.401 | 11.470 | 0.069 | 0.000 | 0.000 | 0.998 | - | - |
| Phk-3 | 21.5 | 8.34E+06 | 3.745 | 3.478 | -0.267 | 0.000 | 0.002 | 0.998 |  | Yes |
| Dmel\CG10211 | 2.9 | 3.08E+05 | 3.546 | 4.039 | 0.493 | 0.000 | 0.000 | 0.998 |  | Yes |
| Hsp67Bc | 23.6 | 2.29E+06 | 3.513 | 4.133 | 0.620 | 0.002 | 0.002 | 0.998 |  | Yes |
| Rac1 | 12.5 | 3.32E+06 | 3.039 | 1.662 | -1.377 | 0.001 | 0.300 | 0.998 | Yes |  |
| BicDR::GFP | 59.9 | 1.85E+07 | 2.804 | 3.021 | 0.217 | 0.002 | 0.024 | 0.998 | - | - |
| l(2)k01209 | 4.6 | 1.11E+06 | 2.670 | 0.707 | -1.963 | 0.015 | 0.774 | 0.998 | Yes |  |
| Kri | 15.3 | 1.25E+07 | 2.242 | 1.486 | -0.756 | 0.017 | 0.103 | 0.998 | Yes |  |
| RpS5a | 49.6 | 2.85E+08 | 2.181 | 1.771 | -0.410 | 0.024 | 0.099 | 0.998 | Yes |  |
| tou | 1.8 | 2.80E+05 | 2.015 | -0.341 | -2.357 | 0.004 | 0.880 | 0.998 | Yes |  |
| Mtl | 10.8 | 1.03E+07 | 1.796 | 1.192 | -0.604 | 0.047 | 0.121 | 0.998 | Yes |  |
| mey | 8.0 | 4.32E+06 | 1.741 | 0.498 | -1.243 | 0.005 | 0.648 | 0.998 | Yes |  |
| Srp54k | 35.0 | 2.57E+07 | 1.706 | 1.563 | -0.143 | 0.004 | 0.024 | 0.998 |  | Yes |
| RpS4 | 63.2 | 2.97E+08 | 1.689 | 1.373 | -0.317 | 0.004 | 0.076 | 0.998 | Yes |  |
| RpL12 | 52.1 | 5.18E+08 | 1.680 | 1.538 | -0.142 | 0.047 | 0.103 | 0.998 | Yes |  |
| RpS5b | 43.5 | 1.11E+09 | 1.529 | 1.406 | -0.123 | 0.013 | 0.036 | 0.998 |  | Yes |
| RpS19a | 96.2 | 3.66E+09 | 1.366 | 1.288 | -0.078 | 0.047 | 0.100 | 0.998 | Yes |  |
| RpL27A | 53.7 | 1.30E+08 | 1.315 | 1.234 | -0.081 | 0.021 | 0.100 | 0.998 | Yes |  |
| RpS17 | 84.0 | 1.53E+09 | 1.290 | 1.210 | -0.079 | 0.016 | 0.103 | 0.998 | Yes |  |
| RpL35 | 35.0 | 2.15E+08 | 1.289 | 0.872 | -0.417 | 0.047 | 0.196 | 0.998 | Yes |  |
| Gp93 | 18.2 | 1.07E+07 | 1.273 | 0.688 | -0.585 | 0.021 | 0.316 | 0.998 | Yes |  |
| RpL13 | 62.8 | 5.56E+08 | 1.143 | 0.826 | -0.317 | 0.047 | 0.205 | 0.998 | Yes |  |
| RpL31 | 79.8 | 3.79E+08 | 1.104 | 0.915 | -0.189 | 0.047 | 0.121 | 0.998 | Yes |  |
| RpS18 | 73.7 | 1.42E+09 | 1.095 | 0.983 | -0.112 | 0.047 | 0.112 | 0.998 | Yes |  |

**Table S2.**
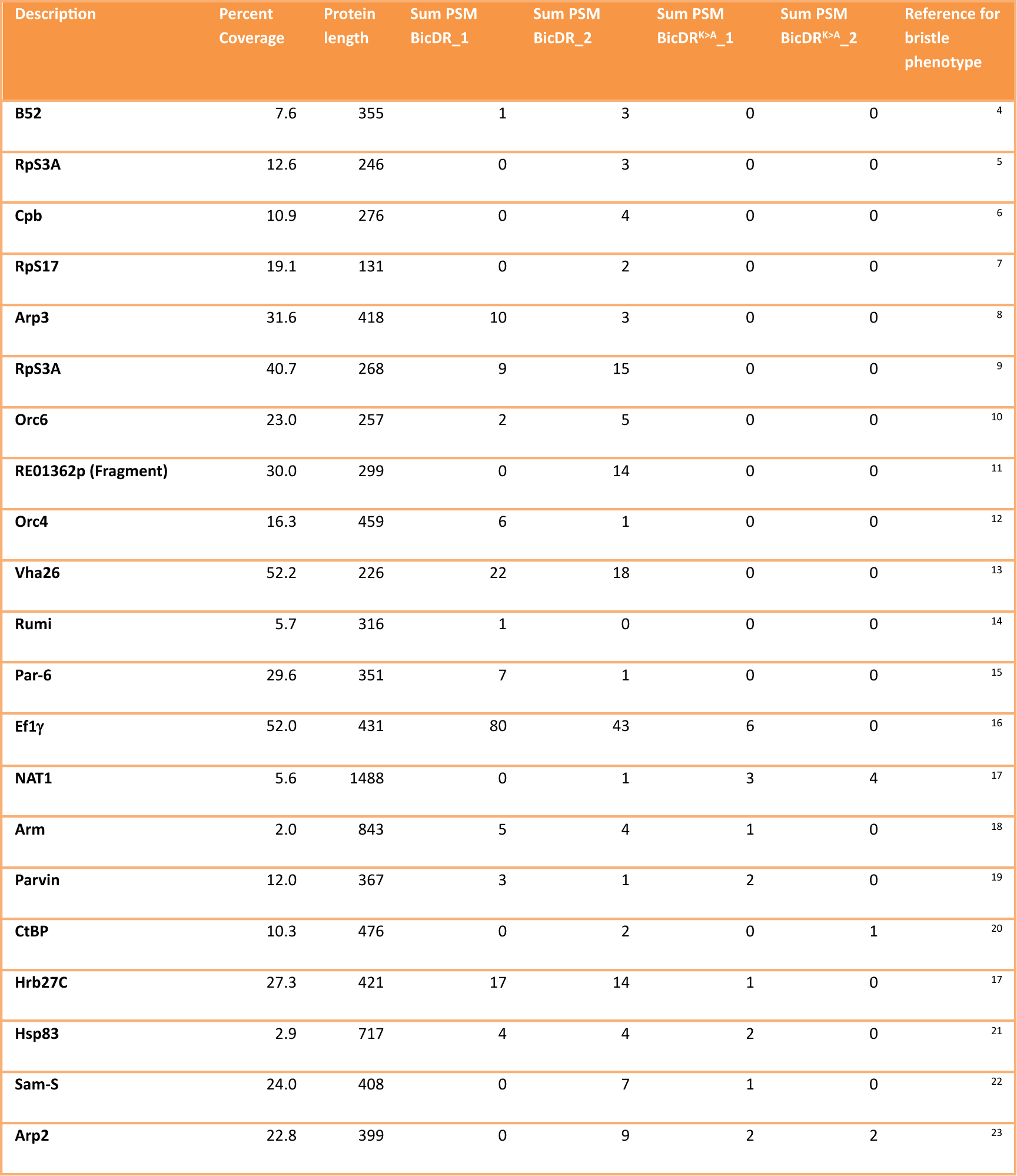
Out of the 118 proteins identified in the SDS-PAGE bands of BicDR::GFP immunoprecipitations, 21 are known to result in bristle phenotypes if perturbed. Abbreviations for BicDR::GFP: BicDR and BicDR^K555A^::GFP: BicDR^K>A^. The Sum PSM describes the summarized number of peptide spectrum matches of the sample with the indicated genotype. Primary data is from Supplementary Data file S2.

